# Knowledge, attitude and practices on dengue fever among paediatric and adult in-patients in Metro Manila, Philippines

**DOI:** 10.1101/520981

**Authors:** Von Ralph Dane M. Herbuela, Ferdinand S. de Guzman, Girly D. Sobrepeña, Andrew Benedict F. Claudio, Angelica Cecilia V. Tomas, Carmina M. Arriola-delos Reyes, Rachele A. Regalado, Mariama M. Teodoro, Kozo Watanabe

## Abstract

**Background:** Knowledge, attitude and practice (KAP) studies have included mainly community-based samples, yet, investigation on in-patients with Dengue fever (DF) through hospital-based surveillance has not been done.

**Methods:** This study aimed to assess and compare the KAP, identify its determinants and protective factors among 250 clinically or serologically confirmed paediatric (n = 233) and adult patients (n = 17) with DF and 250 youth (n = 233) and adult (n = 17) controls.

**Results:** Paediatric patients with DF had significantly higher knowledge (P < 0.05) and practice (P < 0.05) domains mean scores than adult patients with DF and significantly lower practice mean scores than youth controls (P < 0.05). Being senior high school, days in the hospital and rash determined increased KAP among paediatric patients with DF while no significant determinants were found among adult patients with DF. Mosquito-eating fish, screen windows and Dengue vaccine were protective factors against DF, though, further studies should confirm these results. Moreover, there was a significant positive correlation between knowledge and attitude (P < 0.01) of paediatric patients with DF, however, similar with adult patients with DF, these domains did not correlate with their practices against DF.

**Conclusion:** This suggests that the translation of knowledge and attitude to better practices against DF was poor. Thus, it is necessary to structure health programs on models that facilitate behavioural change among children and adults.

## Background

To date, there has been no known cure for Dengue Fever (DF), the world’s fastest spreading mosquito-borne disease which causes approximately 390 million cases per year and puts an estimated 3.9 billion people at risk in 128 countries [1-3]. Since DF epidemiology and ecology are strongly associated with human habits and activities [4], community-based studies have been done to assess the knowledge, attitude and practices (KAP) of people on DF.

Several community-based KAP studies have investigated the correlation among KAP domains. Harapan et al. [5] reported that good knowledge is positively associated with good practice. This is parallel to the report by Alyousefi et al. [6] that poor knowledge on DF has significant positive association with poor preventive practices. However, other similar studies had different results. Kumaran et al. [7] and Shuaib et al. [8] reported that knowledge on causes, signs, symptoms, mode of transmission and preventive practices against DF is not correlated with the practice of preventive measures against DF. Aside from these, two case-control studies reported which preventive practices are protective factors against DF. Regression models revealed that removing trash and stagnant water from around the residence, using mosquito repellent oils, use of mosquito bed nets, fumigation inside the house, and piped water inside the house can reduce the risk and vulnerability to DF infection [9, 10].

Most of the KAP studies have included only community-based samples and investigation on hospital-based samples with clinical or serologically-confirmed DF diagnosis has not been done. Chen et al. [9] interviewed patients who were randomly sampled from a web-based reporting system through telephone interviews. However, this method limits the collection to individuals and households with telephones. It also had 50% response and completion rate among respondents [10, 11]. On the other hand, face to face interview with questionnaire would obtain good response and acceptance rate (99%) and a low refusal rate (1%) among in- and out-patients [11, 12]. Kenneson et al. [10] also did clinical ascertainment and community screening to interview households with and without DF infections by identifying acute or recent DF infections. However, the data collected among households with acute or recent DF infections suggested self-report bias, as members of these households may have already acquired knowledge, changed their behaviour or attitude towards DF during their surveillance [10]. Therefore, hospital-based surveillance, compared with community-based surveillance, would allow us to capture patients’ knowledge and attitude and their family’s/household’s practices against DF during hospitalization (acute phase [febrile-critical] of the infection [2-7 days] from the onset of fever [1]).

Previous KAP studies have also reported that sociodemographic data like income, employment, education, marital status, religion, sex, age, location, socio-economic status, type of residence and DF history were associated with KAP [5, 8, 13-21]. However, to our knowledge, no study has investigated the association between clinical parameters (e.g. diagnosis, platelet count), clinical symptoms (e.g. fever, rash, abdominal pain) and KAP, more so, the difference of determinants of KAP between paediatric and adult patients with DF. Since adults exhibit higher incidence of the severe forms of DF compared with children [22], clinical presentations of symptoms may also be significantly different between paediatric and adult patients with DF. Vomiting and skin rash were more prevalent among children while myalgia and arthralgia, nausea, thrombocytopenia were more exhibited by adult patients with DF [22, 23].

Based on the literatures presented, we hypothesized that paediatric and adult patients’ knowledge and attitude on DF would not have significant positive relationship with their practices against DF, compared with the youth and adult controls. We also hypothesized that paediatric and adult patients with DF may have different determinants of KAP by socio-demographic profile like age, gender and education. In addition, clinical symptoms as determinants of KAP will be different between paediatric and adult patients with DF. Vomiting and skin rash would determine high KAP among paediatric patients with DF while myalgia and arthralgia, nausea, thrombocytopenia would de determinants of high KAP among adult patients with DF. With all these, we hypothesized that paediatric and adult patients with DF would have lower levels of KAP domains than the youth and adult controls, which would also give us hints on which KAP domain have aggravated the acquisition of the infection. Therefore, this study aimed to assess and compare the KAP of paediatric patients and adult patients with DF, paediatric patients with DF and youth control and adult patients with DF and adult controls. We also sought to identify the determinants of KAP domains by socio-demographic profiles, clinical parameters and symptoms, analyse the relationship among the KAP domains, and identify protective factors against DF.

## Methods

### Study and Sampling Design

This study used a non-probability purposive sampling method among patients with DF admitted in 3 public tertiary (>100 beds) hospitals in Metro Manila, Philippines: San Lazaro Hospital, a referral facility for Infectious/ Communicable Diseases, Quezon City General Hospital and; Pasay City General Hospital during the rainy season from 26^th^ July to 26^th^ November 2017. A sample size of 355 was recommended to assume that 50% of patients had good KAP on DF, with a 5% margin of error, 95% CI (a = 0.05; critical value/Z-score of 1.96) based on 4,525 cases in Metro Manila, Philippines, during the same period in 2016 [24]. The number of DF cases increased by 15.5% in Metro Manila from January 1 to May 6 (morbidity week 1–18), which was one of the highest rates in the country in 2017 [25]. For the controls, we followed the 1:1 ratio (one case patient/ one control) with an assumed odds-ratio of ≥2, power (1-β) of 0.80, 0.05 significance level, Z_α_=1.96 [26].

### Participant Inclusion and Exclusion Criteria

A semi-structured bedside interview was done among paediatric (<19 years old) and adult in-patients (age >18 years old) with serology-confirmed or clinically diagnosed DF, who were conscious and able to read and write. Excluded were those who were not able to comply with consent procedures, or with life-threatening comorbidities. Controls were randomly sampled individuals who had no signs and clinical symptoms of DF and who had no family member hospitalized for or diagnosed with DF at the time of interview. Community-based controls were compared with adult patients with DF while paediatric patients with DF were compared (8 to 18 years old) with school-based Grade 3 to Grade 12 students.

### Ethical Considerations

The study was written and conducted based from international and local ethical guidelines: Declaration of Helsinki, ICH-GCP Guidelines and National Ethical Guidelines for Health Research [27-29]. It was reviewed and approved by the Institutional Ethics and Review Boards (IERBs) of each participating hospital: Research Ethics and Review Unit of San Lazaro Hospital, Research Ethics and Technical Committee of Pasay City General Hospital and Planning, Development, Education and Research office of Quezon City General Hospital. Informed consent was obtained from all the controls and patients and/or their parent or legally authorized representative (LAR), or caregiver, especially of those who were under 18 years old.

### Forms and Instruments

Socio-demographic Profile, Clinical Parameters and Symptoms. Both patients and controls were asked about their personal information like age, civil status, gender, educational attainment or employment status, and family monthly income and family and self DF history. Patients’ clinical parameters like admitting diagnosis, serologic test results (NS1Ag and BLOT: IgG and IgM) and laboratory data (*i.e.*, CBC with platelet count) were obtained from medical charts which were used to identify their current DF phase (acute: febrile to critical and recovery phase). Clinical symptoms or chief complaints were also asked.

KAP about DF was developed by Shuaib et al. [8] in Jamaica which was pretested and completed three Delphi Method review rounds for question and response construction and purpose of the questionnaire. The survey has 3 domains: 29-item knowledge (dengue symptoms, modes of transmission, preventive practices and disease management), 3-item attitudes (seriousness, risk and prevention) and 12-item practices (mosquito-man contact and eliminating breeding sites) [8]. Knowledge and attitude domains pertain to each participant’s self-report of knowledge and perception towards DF, while the practice domain involves each participant’s household-report of the preventive practices against DF. We added two items in the list of sources of information (e.g. social media and “barangay” or villages and community) and 1 item in practice (e.g. dengue vaccine). A three-point scale, “yes”, “no” and “I don’t know” was used in knowledge domain. Correct responses were coded 1, otherwise, coded 0 [18]. A 5-point scale, “strongly agree” to “strongly disagree” was used to identify participants’ attitudes where “strongly agree” scored 2 and “agree” scored 1. Likewise, one item in practices (frequency of cleaning ditches and containers with water) used 4-point scale of “always” to “never” where “always”, “often”, “sometimes” were scored 3, 2 and 1, respectively. For more information on the permission acquisition and translation and validation procedures of the questionnaire, please see Additional File 1.

### Statistical and Data Analysis

Statistical analysis was done using Statistical Package for Social Sciences (SPSS) version 25 (IBM Corp., Armonk, NY). We compared the groups: paediatric and adult patients, youth and adult controls, paediatric patients and youth controls, and adult patients and adult controls by their mean scores in each KAP domain using independent samples t-test. To identify determinants (socio-demographic profiles, clinical parameters and clinical symptoms) of KAP, we did a multiple linear regression analysis. It was conducted by inputting socio-demographic and clinical variables (dummy variables [i.e., 0 or 1] for categorical variables) in the model using a stepwise method in backward selection to identify significant (P < 0.05) determinants of KAP. We also calculated the difference in the proportion of participants (yes vs. no) in each source of information using chi-square test. Then, we compared their mean scores in each source of information to identify which increases KAP levels by using independent t-test. To calculate the correlation values between the KAP domain scores, Spearman’s rank correlation (r_s_) (two-tailed) and the fisher’s R-to-Z transformation to obtain confidence interval (CI) were used for the not normally distributed scores as shown by the Shapiro-Wilk and Kolmogorov-Smirnov normality tests [18]. All preventive practices were used in a logistic regression analysis to identify protective factors against DF infection in youth and adult samples. All significant factors (P < 0.05) were put in the multiple regression analysis using stepwise backward selection method.

## Results

### Socio-demographic Profile, Clinical Parameters and Symptoms

Initially, there were 350 patients with DF participated in the study. However, we have excluded those who had incomplete responses (n = 15, 4.3%) and those whose responses came from a family member instead of the patient himself (n = 85, 24.3%). Thus, data from 500 participants comprising of 250 patients with DF (paediatrics n = 233 [93.2%]; adults n = 17 [6.8%]) and 250 controls (youth n = 233; adults n = 17) were included in the final analysis. Paediatric patients with DF had a mean (M) age of 13, and an SD (±) of 3.16 years. All were single (100%), 56.7% were males, and 46.9% were in junior high school, 84% belong to a family with a monthly income of ≤10,000 pesos. The age of adult patients ranges from 19 to 49 years old (M, 29.9±10), 64.7% were females; 73.3% were single, 61.5% were employed, 70.6% belong to a family with a monthly income of ≤10,000 pesos and 70.6% belong to a family with ≤5 members. All (100%) adult patients and majority (77.7%) of paediatric patients with DF had dengue with warning signs. A large proportion of paediatric patients and adult patients had no DF history (92.7% and 93.3%, respectively), had no family DF history (69% and 88.2%, respectively) and were in the acute (febrile-critical) phase of the infection (80.7% and 70.6%, respectively). More than half (68.2%) of paediatric patients and majority (88.2) of adult patients had thrombocytopenia (9,900/mm^3^). Nearly half (43%) of paediatric patients and 35.3% of adult patients had petechiae or rashes.

Furthermore, youth controls had a mean age of 14.11 (±1.88) years with almost half (47.6%) belong to 14-16 age group while adult controls had a mean age of 26.6 (±6.07) years which range from 20 to 46 years old. Half (51.1%) of youth controls were males while 64.7% of adult controls were females. All (100%) of youth controls and majority (94.1%) of adult controls were single. Most of the youth (92.7%) and adult controls (93.8%) belong to a family with a monthly income of ≥10,000 pesos. There was a preponderance of youth (86.7%) and adult (94.1%) controls who had no DF history. More than half of youth (79.4%) and adult controls (58.8%) had no family DF history. For more information on the profile of patients with DF and controls, please see Additional file 2: Table S1.

### Prevalence of good knowledge, attitudes and practices

An independent samples t-test revealed that paediatric patients (PP) obtained significantly higher mean scores than adult patients (AP) in knowledge (P = 0.03) and practice (P = 0.02) domains as shown in **Figure 1.** In control group, adult controls (AC) had significantly higher mean scores than youth controls (YC) in all domains: knowledge (P < 0.001), attitude (P < 0.001) and practice (P = 0.02). When we compared the mean scores of KAP domains between paediatric patients with DF and youth controls, the former had significantly higher mean scores than the latter in knowledge (P < 0.001) and attitude (P < 0.001) domains. As expected, paediatric patients with DF obtained lower mean score than youth controls in practice domain (P = 0.03) and adult patients with DF had significantly lower mean scores than adult controls in knowledge (P < 0.001) and practice (P < 0.001) domains.

**Fig. 1.** Independent t-test results for the difference of knowledge, attitude and practices domains mean scores among paediatric and adult patients with DF and youth and adult controls. PP = Paediatric Patients, AP = Adult Patients, YC = Youth Controls, AC = Adult Controls; *p < 0.05; **p < 0.01; ***p < 0.001

### Determinants of knowledge, attitudes and practices

Multiple linear regression analysis found significant regression equations in all KAP domains among paediatric patients with DF as shown in **Table 1**. It showed that knowledge increased significantly more in paediatric patients with DF who were senior high school while it decreased significantly more in paediatric patients who were in college and those who had DF for the first time. Being senior high school also tended to increase paediatric patients’ attitude. Then, as their days in the hospital increases, their attitude scores also increase, however, as their age increases, their attitude score decreases. Further, practice scores tend to decrease among those with severe dengue, however, it tends to increase to those paediatric patients who had petechiae or rash. Age was found to increase knowledge, attitude and practice, being female increased both knowledge and attitude and having family DF history increased attitude among youth controls. While no significant determinants were found among adult patients with DF, being in college or university, being female and being in a family with more than 5 members decreased attitude and being unemployed and having DF for the first time, decreased practice among adult controls.

**Table 1.** Multiple linear regression results showing the determinants of KAP among patients with DF and controls

| Outcome Variables | Determinants |  | R<sup>2</sup> | β | p-value |
| --- | --- | --- | --- | --- | --- |
| Knowledge | Paediatric patients | Education (senior high school) | 0.09 | 0.18 | 0.01 |
|  |  | Education (college) |  | -0.16 | 0.02 |
|  |  | No DF history (first-time) |  | -0.15 | 0.04 |
|  | Youth controls | Age | 0.18 | 0.38 | < 0.001 |
|  |  | Female |  | 0.19 | 0.002 |
| Attitude | Paediatric patients | Age | 0.11 | -0.30 | 0.004 |
|  |  | Education (senior high school) |  | 0.39 | < 0.001 |
|  |  | Days in the hospital |  | 0.16 | 0.04 |
|  | Youth controls | Age | 0.09 | 0.19 | 0.003 |
|  |  | Female |  | 0.16 | 0.01 |
|  |  | Had family DF history |  | 0.14 | 0.03 |
|  | Adult Controls | Education (college) | 0.63 | -0.48 | 0.01 |
|  |  | Female |  | -0.48 | 0.01 |
|  |  | Family members (> 5) |  | -0.38 | 0.04 |
| Practice | Paediatric patients | Severe dengue | 0.07 | -0.15 | 0.04 |
|  |  | Rash or petechiae |  | 0.18 | 0.01 |
|  | Youth controls | Age | 0.06 | 0.24 | < 0.001 |
|  | Adult controls | Unemployed | 0.57 | -0.92 | < 0.001 |
|  |  | No DF history (first-time) |  | -0.58 | 0.008 |

β = standardized beta coefficients; DF = dengue fever

### Correlation among knowledge, attitudes and practices

Spearman rank correlation revealed that there was a significant positive correlation between knowledge and attitude domains of paediatric patients with DF as shown in **Table 2**. However, as hypothesized, there was no correlation found in knowledge-practice and attitude-practice domains of both paediatric and adult patients with DF. Among controls, only youth controls had obtained significant positive correlations among the KAP domains, wherein a strong correlation was found between knowledge-practice domains with a correlation coefficient of 0.42 (95% CI: 0.34-0.57).

**Table 2.** Correlation between the KAP domains among patients with DF and controls

| Patients with DF |  |  | Controls |  |  |
| --- | --- | --- | --- | --- | --- |
| Variables | $r_s$ (95%CI) | *p*-value | | $r_s$ (95%CI) | *p*-value |
| **Knowledge-attitude** |  |  |  |  |  |
| Paediatric | 0.20 (0.08, 0.13) | 0.002 | Youth | 0.34 (0.24-0.48) | < 0.001 |
| Adult | 0.02 (-0.51, 0.59) | 0.95 | Adult | -0.05 (-0.50, 0.60) | 0.84 |
| **Knowledge-practice** |  |  |  |  |  |
| Paediatric | 0.06 (-0.06, 0.20) | 0.36 | Youth | 0.42 (0.34-0.57) | < 0.001 |
| Adult | -0.39 (-0.81, 0.24) | 0.12 | Adult | 0.03 (-0.58, 0.52) | 0.91 |
| **Attitude-practice** |  |  |  |  |  |
| Paediatric | -0.04 (-0.15, 0.10) | 0.57 | Youth | 0.23 (0.13-0.38) | < 0.001 |
| Adult | -0.13 (-0.62, 0.48) | 0.62 | Adult | 0.19 (-0.41, 0.68) | 0.46 |

$r_s$ : Spearman rank correlation coefficients; 95% Confidence intervals (CI) were transformed using Fisher's R-to-Z

### Sources of information on DF

Television (TV) was the main source of information among patients with DF (75.2%) and controls (72%). Chi-square test analyses showed that paediatric and adult patients with DF were more likely to get information on DF from hospital, doctors and nurses (68.8%, P < 0.001) and health centres (64%, P < 0.001), compared with youth and adult controls. More than half of youth (60.5%) and adult controls (53%) identified social media (e.g. Facebook, Twitter, Instagram etc.) (60 %, P < 0.001) and family members (63% and 59%, P < 0.001) and school (55.8% and 52.9%, P 0.04) as their sources of information about DF. Further analysis of mean score comparisons using independent t-test found that paediatric and adult patients who reported to have obtained information on DF through newspaper and health centre, respectively, had higher knowledge mean scores. In youth controls, those who have obtained information on DF through social media, newspaper, health brochures, family members, school, hospital, doctors and nurses and health centres had higher mean scores in knowledge domain. Attitude and practice mean scores were higher among those paediatric patients with DF and youth controls who had identified newspaper, health brochures, family, school, hospital, doctors and nurses, barangay and community and health centres as their sources of information on DF. Adult controls who reported neighbours as their source of information on DF also had high attitude mean scores and those who had high practice mean scores reported barangay and community and workplace as their sources of information on DF. (Additional file 3: Table S2).

### Practices against DF

All preventive practices were used in a logistic regression analysis to identify protective factors against DF. Then, after a multivariate regression analysis, use of mosquito eating fish, Dengue vaccine, use of screen windows, and doing at least one preventive practice against DF were found to be protective factors against DF among youth samples (paediatric patients with DF and youth controls) as shown in **Table 3**. Among adults (adult patients with DF and adult controls), only the use of screen windows was identified as a significant protective factor against DF with adjusted odds-ratio (aOR) of 23.9 (95% CI: 2.08-275.2, P = 0.01). For both youth and adult samples, mosquito eating fish, screen windows, and Dengue vaccine were identified as protective factors against DF infection. The strongest factor in the model was use of mosquito eating fish, with an adjusted ratio (aOR) of 8.69 (95% CI: 3.67-20.57, P = < 0.001).

**Table 3.** Multiple logistic regression model of predictors of absence of DF infection

| Likelihood Ratio Estimates |  |  |  |  |  |  |
| --- | --- | --- | --- | --- | --- | --- |
| Practices | DF | $\beta$ | SE | Wald $\chi^2$ | aOR (95% CI) | p-value |
| **Youth** |  |  |  |  |  |  |
| screen windows | 1 | 1.45 | 0.31 | 21.15 | 4.25 (2.29-7.88) | < 0.001 |
| eliminate standing water | 1 | -1.40 | 0.41 | 11.96 | 0.24 (0.11-0.54) | 0.001 |
| mosquito eating fish | 1 | 2.03 | 0.45 | 20.70 | 7.62 (3.18-18.3) | < 0.001 |
| does nothing to reduce mosquitoes | 1 | 0.98 | 0.37 | 6.87 | 2.67 (1.28-5.57) | 0.009 |
| Dengue vaccine | 1 | 1.60 | 0.28 | 32.58 | 4.95 (2.86-8.56) | < 0.001 |
| covering water containers | 1 | -2.32 | 0.53 | 19.15 | 0.10 (0.03-0.28) | < 0.001 |
| **Adults** |  |  |  |  |  |  |
| professional pest control | 1 | 1.82 | 0.93 | 3.88 | 6.20 (1.01-38.1) | 0.05 |
| screen windows | 1 | 3.17 | 1.25 | 6.49 | 23.9 (2.08-275) | 0.01 |
| **Both** |  |  |  |  |  |  |
| screen windows | 1 | 1.53 | 0.30 | 25.92 | 4.60 (2.56-8.28) | < 0.001 |
| eliminate standing water | 1 | -1.41 | 0.39 | 13.20 | 0.24 (0.11-0.52) | < 0.001 |
| mosquito-eating fish | 1 | 2.16 | 0.44 | 24.23 | 8.69 (3.67-20.6) | < 0.001 |
| does nothing to reduce mosquitoes | 1 | 0.70 | 0.36 | 3.83 | 2.01 (1.00-4.04) | 0.05 |
| Dengue vaccine | 1 | 1.49 | 0.27 | 30.32 | 4.42 (2.61-7.55) | < 0.001 |
| covered water containers | 1 | -2.01 | 0.49 | 17.09 | 0.13 (0.05-0.35) | < 0.001 |

DF = degree of freedom; $\beta$ = standardized bet coefficients; Wald $\chi^2$ = Wald chi-square; aOR (95% CI) = adjusted odds-ratio 95% confidence interval

## Discussion

There was a positive correlation found between knowledge and attitude domains of paediatric patients with DF. These indicate that there was a good translation of knowledge to attitude on DF among paediatric patients with DF. Their knowledge on dengue symptoms, modes of transmission, preventive practices against DF and disease management tend to have changed their beliefs that DF is a serious and threatening disease. Although paediatric patients’ knowledge correlated with their attitude towards it, both knowledge and attitude did not correlate with their practices against DF. These findings clearly signify that the translation of knowledge and attitude to practice among paediatric patients were poor. This was also found to be true among adult patients with DF, their knowledge and attitude on DF was not correlated with their practices as well. This means that although paediatric and adult patients with DF were knowledgeable about the symptoms of DF, vector breeding sites control, transmission modes of DF, and perceived DF as a serious and threatening disease, it did not lead to change in their behaviour of doing the preventive practices against it. This implies that the poor practice against DF might have exposed them to higher risk of contracting the disease.

The results suggest that health programs should be designed for children and adolescents which focus on translating their knowledge and attitudes into better and effective practices against DF through behaviour change. Many programs continue to focus only on changing people’s knowledge and on raising awareness, rather than physical activity programs which are more successful at producing behaviour change [30]. The Communication for Behavioural Impact (COMBI), a comprehensive strategy that uses communication for knowledge to have significant effect to behavioural change (making people becoming aware, informed, convinced, and deciding to act, then repeating and maintaining that action) or increased practices against DF [7,31]. Moreover, another model that facilitates behavioural change that could increase the translation of attitude to practice among children and adolescents is the Health Belief Model (HBM). This model suggests that a change in the behaviour or acting can be expected if a person perceives themselves to be at risk or susceptible to the disease (perceived susceptibility), that the disease will have serious consequences (perceived severity), a course of action will minimize consequences (perceived benefits), and the benefits of action will outweigh the cost of barriers (perceived barriers) and self-efficacy [32]. Both models should on changing the behaviour not only in individual and household levels but also in community level because community participation, including schools, especially children, is necessary to effectively control the vector mosquitoes [33].

Previous studies have reported that there is a significant positive correlation between education and level of knowledge and attitude toward DF [17, 21]. However, in our study, paediatric patients who were senior high school tend to have increased knowledge and attitude on DF, compared to those in college or university. One possible reason could be senior high schools may have included contents about DF in their curriculum that may have increased the knowledge and attitude levels of paediatric patients in senior high school. Having DF history was also found to be a significant determinant of high knowledge mean scores among paediatric patients with DF. It may appear obvious; however, past experiences such as infection, which is prevalent among children [28] increased paediatric patients’ knowledge about it. Moreover, as expected, age was found to increase knowledge, attitude and practices among youth controls, however, in attitude domain, the opposite was found among paediatric patients with DF. As the age of paediatric patients with DF increases, their attitude towards DF decreases. Hospitalized younger children were reported to be very conscious about their health and express very positive health attitudes than older children [34,35]. This may be brought about by the fact that younger children who are hospitalized are more vulnerable to emotional upset and they experience greater anxieties, arising from separation from parents [36, 37].

Another significant determinant of attitude towards DF was the number of days in the hospital. Paediatric patients who identified hospital, doctor and nurses (70%) and health centres (65%) as their sources of information on DF, had significantly higher attitude mean scores than those who didn’t identify these as their sources of information on DF. This implies that the longer they stay in the hospital, the more they perceive DF as serious and threatening which may be due to their experience of anxiety towards medical settings and receiving of medical care [38], especially fear of medical procedures like injection needles [39] (daily drawing of blood to check their platelet counts) and they perceive medical professionals like doctors and nurses as inflictors of trauma [40]. Aside from these, hospitalization also increases the chance of children to be dissatisfied with their hospital-stay situations like food conditions [41]. Paediatric patients with DF were advised to avoid eating dark coloured foods (to monitor the colour of stool for signs of bleeding) [42]. Hospitalized children also experience anxiety because of limited physical activities like being absent in school, limited chance of spending time and playing with peers and friends or siblings [43].

Petechiae or rash was found to be a significant determinant of attitude among adult patients with DF. As one of the common (35.3%) reported symptoms among adult patients with DF in this study, this implies that those who had petechiae or rashes were more likely to gain belief that DF is a serious illness and anyone is at risk of it. These cause itching and swelling of the palms/soles [44] and its presence may signify severity of the disease as it is the most seen and observed among the symptoms which would have triggered the high attitude level toward DF among adult patients. One possible reason why petechiae or rash was a significant determinant of attitude among adult patients with DF, and not among paediatric patients with DF, was its high prevalence among paediatric patients with DF, compared with adult patients with DF [23]. Thus, the presence of this symptom didn’t affect the perception of paediatric patients toward DF, instead, the time when it appeared may explain, in partial, why it was found a significant determinant of practices among paediatric patients with DF. The presence of petechial rash (which is also described as “isles of white in the sea of red”) and pruritus (severe itching of the skin) occur towards the end of acute (febrile) phase and the beginning of the recovery phase [1]. This could mean that those paediatric patients who were already having rashes during the interview may have already changed their family/household members’ behaviour and started doing the preventive practices against DF, thus, higher practice mean scores. Severe dengue was also a significant determinant of decreased practice among paediatric patients with DF. Paediatric patients with severe dengue, compared to those who had other DF diagnoses, had the significantly lowest mean score in practice domain. 50% of paediatric patients with severe dengue had to be confined in the intensive care unit (ICU). They were interviewed only after ICU confinement which was, on average, the 5^th^ day of hospitalization. The time spent in the ICU might have decreased the opportunity of their family/household members to immediately do the practices against DF, thus, lower practice mean score.

The difference between paediatric patients with DF and youth controls can also be seen in their sources of information. Youth controls were more likely to get more information on DF from their family members (63%), social media (e.g. facebook, Instagram, twitter, etc.) (61.3%) and school (56%) compared with paediatric patients with DF. Further analysis showed that youth controls, compared to paediatric patients with DF, have obtained more knowledge on DF through the use of social media, newspaper, health brochures, family members, school and hospital, doctors or nurses. This hints that unlike paediatric patients with DF, youth controls had more access to different sources of information on DF which might have improved their knowledge on DF. For example, social media and the use of smartphones, have been seen to be a novel system for disease epidemiology [45, 46]. It has high acceptance rate among younger population who perceived that it could be an effective strategic health communication effort to raise dengue-related concerns in the future [47]. Newspaper, health brochures, family, hospital, doctors and nurses and school were found to have improved youth controls’ knowledge and practice against DF which suggests that these might be effective means to increase not only the knowledge, but also the practices against DF of children.

Multiple logistic regression analysis on the practices revealed that use of mosquito-eating fish, use of screen windows and Dengue vaccine were protective factors against DF. Studies have found that larvivorous fish (*Gambusia Affinis*), the common guppy (*Poecilia reticulata*), *Cyprinidae* or Tilapia spp. can be effectively used to control the mosquito population at their larval stages [48-50]. However, our data collected from both patients with DF and controls, through the use of questionnaire is subjective which might have produced false positive responses, in this case, use of mosquito-eating fish. We had no means to confirm if the participants, especially the patients really had mosquito-eating fish at home, thus, future studies should include direct household observation to validate this result. Moreover, in our study, we assumed that the use of screen windows equates to the use of glass windows in airconditioned rooms among controls, especially the youth. Screen and glass windows could be potential ways to reduce DF transmission by the reduced exposures to vectors that enter homes through open windows [9]. These may not have been available to the patients with DF because majority (85.2%) of them belong to households with low monthly family income, thus, increasing their vulnerability to DF infection. However, our data collected from both patients with DF and controls, through the use of questionnaire is subjective which might have produced false positive responses, which we consider one of the limitations of this study. We had no means to confirm if the participants, especially the patients with DF really had mosquito-eating fish at home or had been using screen windows, thus, future studies should include direct household observation to validate these results.

Surprisingly, Dengue vaccine was found to be a protective factor against DF among youth samples. Another limitation of this study was, we couldn’t rely to the participants’ responses about their Dengue vaccine acquisition history. We had no means to confirm whether they got vaccinated or not. Thus, this result may require more intensive studies to whether it could truly be a protective factor against DF infection. The WHO issued a conditional recommendation in April 2016 on the use of the vaccine for highly dengue-endemic areas due to a subset of trial participants who were inferred to be seronegative at time of first vaccination had a higher risk of more severe dengue and hospitalizations from dengue compared to unvaccinated participants. They still recommend preventive practices that combat vector mosquitoes to control and prevent transmission of DF infection [2].

## Conclusions

Paediatric patients with DF had significantly higher mean scores in knowledge and attitude than youth controls, who, in turn, had significantly higher mean score in practice domain compared with paediatric patients with DF. Being senior high school, days in the hospital and rash or petechiae determined increased knowledge, attitude and practices, respectively, among paediatric patients with DF. There was a significant positive correlation between knowledge and attitude of paediatric patients with DF while their knowledge and attitude were not correlated with their practices against DF. These suggest that although paediatric patients had high knowledge and attitude on DF, its translation to better practice of preventive measures against DF was poor compared with youth controls. These findings highlight the importance of behavioural change for knowledge and attitude to have significant effect to practices against DF among children through health programs campaign which are structured from COMBI and HBM. This study also adds to the emerging topics on protective factors against DF, such as use of mosquito-eating fish, use of screen windows and Dengue vaccine, however, further studies are needed to confirm these results.

To our knowledge, our study is the first to use hospital-based surveillance that investigated the association of clinical data to KAP domains and described the difference of KAP on DF between paediatric patients and adult patients with DF. This is also the first study to use clinical ascertainment through hospital-based surveillance among paediatric and adult patients with DF. Additionally, there was a high response rate (100%) among patients with DF and various dengue diagnoses among paediatric patients with DF at the three tertiary hospitals, allowing an increased generalizability of study findings. One of this study’s major limitations was the relatively small sample size of adult patients with DF which limit the generalizability of study findings in this population. Moreover, participating hospitals were public tertiary hospitals, where most patients belong to low-income families. Thus, association of income with the domains was hard to estimate. Finally, only in-patients were included in this study, limiting the analysis to those admitted to hospitals. Therefore, we recommend that future studies also include out-patients to see whether hospitalization is confounding the association between the constructs and dengue infection.

## Abbreviations

DF: Dengue fever;
KAP: knowledge, attitude and practices;
ICH-GCP: International Conference on Harmonization-Good Clinical Practice;
LARs: Legally authorized representatives;
NS1Ag: Non-structural protein 1 antigen;
IgG: Immunoglobin G;
IgM: Immunoglobin M;
CBC: complete blood count;
CI: Confidence interval;
M: Mean;
aOR: Adjusted odds ratio;
COMBI: Communication for behavioral impact;
HMB: Health belief model;
WHO: World health organization

## Declarations

### Ethics approval and consent to participate

This study underwent review and was approved by the different institutional ethics research boards (IERBs) of the three hospitals participated in the study: Research Ethics and Review Unit of San Lazaro Hospital, Research Ethics and Technical Committee of Pasay City General Hospital and Planning, Development, Education and Research office of Quezon City General Hospital. Informed consent was obtained from all the patients and their parents, legally authorized representatives (LARs) or caregivers, especially for those who were under 18 years of age and youth and adult controls.

## Consent for publication

Not applicable.

## Availability of data and materials

The datasets supporting the conclusions of this article are included within the article and its additional files. MS_KAP_AF1.docx Additional file 1: Detailed description of the knowledge, attitude and practices (KAP) on Dengue Fever questionnaire permission acquisition, translation and validation procedures (DOCX 16 KB) MS_KAP_AF2.docx Additional file 2: Profile of pediatric and adult patients with DF and youth and adult controls (DOCX 25 KB)

MS_KAP_AF3.docx Additional file 3: Detailed information on the sources of information on dengue fever of pediatric and adult patients with DF and controls. (DOCX 41 KB)

## Author Contributions

Author V.H. designed the study, wrote the protocol, conducted the interviews, analysed the data and written this manuscript. Author F.D.G., G.S. & A.C. were assigned as co-investigators in each hospital site and supervised patient recruitment and data gathering. Author A.T. & C.R. provided guidance and comments on the initial drafts of the study protocol including literature review, sampling, data gathering methods and ethical considerations. Author R.R. supervised the interviews, testing and scoring procedures for the controls. Author M.T. worked with V.H. from the submission to approval of study protocol in the hospitals and over-all data gathering procedures. Author K.W. supervised the data gathering and provided guidance and comments on the analysis and the initial drafts of this manuscript. All authors have contributed to and have approved the final manuscript.

## Acknowledgments

The authors would like to thank Thaddeus M. Carvajal, John Robert R. Bautista, Howell T. Ho for their assistance in the validation of the KAP questionnaire. We would also like to thank Jazteene Dale M. Villarama, Joenna Mari P. Cabarloc, Jayne Nicholei C. Borricano and Ma. Lourdes SJ. Orbeta for their assistance during data gathering. Most importantly, we are grateful to all the patients and controls who participated in this study. This study was supported by the Japan Society for the Promotion of Science (JSPS) Grant-in-Aid for Scientific Research (16H05750, 17H01624) and JSPS Bilateral Joint Research Projects which had no role in the design, data collection, statistical analysis and writing of this manuscript.

## References

1. World Health Organization. Dengue: guidelines for diagnosis, treatment, prevention and control. New edition 2009. http://www.who.int/tdr/publications/documents/dengue-diagnosis.pdf. Accessed 03 May 2017.

2. World Health Organization. Dengue and severe dengue. World Health Organization, 2018. https://www.who.int/news-room/fact-sheets/detail/dengue-and-severe-dengue. Accessed 12 May 2017.

3. Brady OJ, Gething PW, Bhatt S, Messina JP, Brownstein JS, Hoen AG, et al. Refining the global spatial limits of dengue virus transmission by evidence-based consensus. PLoS Negl. Trop. Dis. 2012;6:e1760

4. Yboa BC & Labrague LJ. Dengue Knowledge and Preventive practices among Rural Residents in Samar Province, Philippines. Am J Public Health Res. 2013;1:47–52

5. Harapan H, Rajamoorthy Y, Anwar S, Bustamam A, Radiansyah A, et al. Knowledge, attitude, and practice regarding dengue virus infection among inhabitants of Aceh, Indonesia: a cross-sectional study. BMC Infect Dis. 2018;18:96

6. Alyousefi TA, Abdul-Ghani R, Mahdy MA, Al-Eryani SM, Raja YA, et al. A household-based survey of knowledge, attitudes and practices towards dengue fever among local urban communities in Taiz Governorate, Yemen. BMC Infect Dis. 2016;16:543

7. Kumaran E, Doum D, Keo V, Sokha L, Sam B, et al. Dengue knowledge, attitudes and practices and their impact on community-based vector control in rural Cambodia. PLoS Negl. Trop. Dis. 2018;12:e0006268

8. Shuaib F, Todd D, Campbell-Stennett D, Ehiri J, Jolly PE. Knowledge, attitudes and practices regarding dengue infection in Westmoreland, Jamaica. West Indian Med J. 2010;59:139–46

9. Chen B, Yang J, Luo L, Yang Z & Liu Q. Who is vulnerable to dengue fever? A community survey of the 2014 outbreak in Guangzhou, China. Int J Environ Res Public Health. 2016;13:712

10. Kenneson A, Beltran-Ayala E, Borbor-Cordova MJ, Polhemus ME, Ryan SJ, et al. Social-ecological factors and preventive actions decrease the risk of dengue infection at the household-level: Results from a prospective dengue surveillance study in Machala, Ecuador. PLoS Negl. Trop. Dis. 2017;11:e0006150

11. Safdar N, Abbo LM, Knobloch MJ & Seo SK. Research methods in healthcare epidemiology: Survey and qualitative research. Infect Control Hosp Epidemiol. 2016;37:1272–1277

12. Salkovskis PM, Storer D, Atha C, Warwick HMC. Psychiatric morbidity in an accident and emergency department: characteristics of patients at presentation and one-month follow-up. Br J Psychiatry. 1990;156: 483–487

13. Wong LP, Shakir SM, Atefi N & AbuBakar S. Factors affecting dengue prevention practices: nationwide survey of the Malaysian public. PLoS One. 2015;10:e0122890

14. Syed M, Saleem T, Syeda UR, Habib M, Zahid R, et al. Knowledge, attitudes and practices regarding dengue fever among adults of high and low socioeconomic groups. J Pak Med Assoc. 2010;60:243

15. García-Betancourt T, Higuera-Mendieta DR, González-Uribe C, Cortés S & Quintero J. Understanding water storage practices of urban residents of an endemic dengue area in Colombia: Perceptions, rationale and socio-demographic characteristics. PLoS One. 2015;10:e0129054

16. Paz-Soldán VA, Morrison AC, Cordova Lopez JJ, Lenhart A, Scott TW, et al. Dengue knowledge and preventive practices in Iquitos, Peru. Am J Trop Med Hyg. 2015;93:1330–1337

17. Alves AC, Fabbro AL, Passos ADC, Carniero AFTM, Jorge TM, et al. Knowledge and practices related to dengue and its vector: a community-based study from Southeast Brazil. Rev Soc Bras Med Trop. 2016;49:222–226

18. Dhimal M, Aryal KK, Dhimal ML, Gautam I, Singh SP, et al. Knowledge, attitude and practice regarding dengue fever among the healthy population of highland and lowland communities in central Nepal. PLoS One. 2014;9:e102028

19. Saied KG, Al-Taiar A, Altaire A, Alqadsi A, Alariqi EF et al. Knowledge, attitude and preventive practices regarding dengue fever in rural areas of Yemen. Int Health. 2015;7:420–425

20. Chanyasanha C, Guruge GR & Sujirarat D. Factors influencing preventive behaviors for dengue infection among housewives in Colombo, Sri Lanka. Asia Pac J Public Health. 2015;27:96–104

21. Al-Dubai SAR, Ganasegeran K, Alwan MR, Alshagga MA, Saif-Ali R. Factors affecting dengue fever knowledge, attitudes and practices among selected urban, semi-urban and rural communities in Malaysia. Southeast Asian J Trop Med Public Health. 2013;44:37

22. Namvongsa V, Sirivichayakul C, Songsithichok S, Chanthavanich P, Chokejindachai W, et al. Difference in clinical features between children and adult with dengue hemorrhagic shock syndrome. Southeast Asian J Trop Med Public Health. 2013;44:72–9

23. Souza LJ de, Pessanha LB, Mansur LC, Souza LA de, Ribeiro MBT, et al. Comparison of clinical and laboratory characteristics between children and adults with dengue. Braz J Infect Dis. 2013;17:27–31

24. Department of Health. Weekly Dengue Cases Report, Morbidity Week 28: July 10 – 16 July 2016. Epidemiology Bureau, Public Health Surveillance Division. 2016. https://www.doh.gov.ph/sites/default/files/statistics/DENGUE%20MW28.pdf. Accessed 05 May 2017.

25. Department of Health. Weekly Dengue Cases Report, Morbidity Week 18: January 1 – 6 May 2017. Epidemiology Bureau, Public Health Surveillance Division. 2017. https://www.doh.gov.ph/sites/default/files/statistics/2017_Dengue_MW1-MW18.pdf. Accessed 05 May 2017.

26. Charan J & Biswas T. How to calculate sample size for different study designs in medical research? Indian J Psychol Med. 2013;35:121–126

27. World Medical Association. World Medical Association Declaration of Helsinki Ethical Principles for Medical Research Involving Human Subjects. JAMA. 2013;310:2191–2194.

28. European Medicines Agency. ICH Topic E6 (R1) Guideline for good clinical practice step 5 note for guidance on good clinical practice. (CPMP/ICH/135/95). 2002.

29. Philippine Health Research Ethics Board. National Ethical Guidelines for Health Research. Taguig City: PNHRS; 2011.

30. Beckman H, Hawley S & Bishop T. Application of theory-based health behavior change techniques to the prevention of obesity in children. J Pediatr Nurs. 20016;21:266–275

31. Parks W & Lloyd L. Planning social mobilization and communication for dengue fever prevention and control: a step-by-step guide. Geneva, Switzerland; WHO; 2004

32. Korin MR. Theory and fundamentals of health promotion for children and adolescents. In: Korin M. ed. Health promotion for children and adolescents. Springer, Boston, MA; 2016

33. Elder J & Lloyd L. Working paper 7.3: Achieving behaviour change for dengue control: methods, scaling-up and sustainability. In: Scientific Working Group Report on Dengue: Meeting Report, 1-5 October 2006. Geneva, Switzerland: WHO; 2007

34. Piko BF & Bak J. Children’s perceptions of health and illness: images and lay concepts in preadolescence. Health Educ Res. 2006;21:643–653

35. Woods SE, Springett J, Porcellato L & Dugdill L. ’Stop it, it’s bad for you and me’: experiences of and views on passive smoking among primary-school children in Liverpool. Health Educ Res. 2005;20:645–55

36. Bonn M. The effects of hospitalization on children: a review. Curationis 1994;17:20–24

37. Coyne I. Children’s experiences of hospitalization. J Child Health Care. 2006;10:326–36

38. Wolraich M, Felice ME, Drotar D. The classification of child and adolescent mental diagnoses in primary care: diagnostic and statistical manual for primary care (DSM-PC) child and adolescent version. Elk Grove Village, IL.: American Academy of Pediatrics; 1996.

39. Diaz-Caneja A, Gledhill J, Weaver T, Nadel S, Garralda E. A child’s admission to hospital: a qualitative study examining the experiences of parents. Intensive Care Med. 2005;31:1248–54

40. Pao M & Bosk A. Anxiety in medically ill children and adolescents. Depress Anxiety. 2011;28:40–49

41. Bsiri-Moghaddam K, Basiri-Moghaddam M, Sadeghmoghaddam L & Ahmadi F. The concept of hospitalization of children from the view point of parents and children. Iran J Pediatr. 2011;21:201–8

42. Ong WT. Deadly dengue: Prevention, treatment, and ‘Tawa Tawa’. PCHRD;2014

43. Angström-Brännström C, Norberg A, Jansson L. Narratives of children with chronic illness about being comforted. J Pediatr Nurs. 2008;23:310–316

44. Huang HW, Tseng HC, Lee CH, Chuang HY & Lin SH. Clinical significance of skin rash in dengue fever: A focus on discomfort, complications, and disease outcome. Asian Pac J Trop Med. 2016;9:713–718

45. Roche B, Gaillard B, Leger L, Pelagie-Moutenda R, Sochacki T, et al. An ecological and digital epidemiology analysis on the role of human behavior on the 2014 Chikungunya outbreak in Martinique. Sci. Rep. 2017;7: 1–8

46. Nguyen QC, Brunisholz KD, Yu W, McCullough M, Hanson HA, et al. Twitter-derived neighborhood characteristics associated with obesity and diabetes. Sci. Rep. 2017;7:1–10

47. Lwin MO, Vijaykumaor S, Foo S, Fernand ON, Lim G, et al. Social media-based civic engagement solutions for dengue prevention in Sri Lanka: results of receptivity assessment. Health Educ Res. 2016;31:1–11

48. Noreen M, Arijo AG, Ahmad L, Sethar A, Leghari MF, et al. Biological control of mosquito larvae using edible fish. Int. J. of Inov. and App. Res. 2017;5:1–6

49. Phukon HK & Biswas SP. An investigation on larvicidal efficacy of some indigenous fish species of Assam, India. Adv. Biores. 2013;4:22–25

50. World Health Organization. Comprehensive guidelines for prevention and control of dengue and dengue haemorrhagic fever. WHO, Regional Office for South-East Asia; 2011 http://www.searo.who.int/entity/vector_borne_tropical_diseases/documents/SEAROTPS60/en/

