## Additional File 1 for "Knowledge, attitude and practices on dengue fever among paediatric and adult in-patients in Metro Manila, Philippines"

**Additional file 1 – Detailed description of the knowledge, attitude and practices (KAP) on Dengue Fever questionnaire permission acquisition, translation and validation procedures**

It was first used in a study done in Westmoreland, Jamaica and was approved for use in this study (with modifications) by one of its authors, Dr. Pauline Jolly from the Department of Epidemiology, School of Public Health, University of Alabama at Birmingham, USA. It was translated to Filipino (Tagalog) by independent translators from the *Sentro ng Wikang Filipino* (SWF) (Centre for the Filipino Language) of the University of the Philippines-Manila, Philippines. It was then face, content and construct validated by Public Health and Infectious Disease experts using the Survey/Interview Validation for Expert Panel (VREP)^1^. Then it was administered on a selected hospital and school-based samples. When a consensus was reached, it was translated back to English by bilingual translators from the *Komisyon sa Wikang Filipino* (KWF) (Commission on the Filipino Language). Initially, the translated questionnaire in Filipino was tested for internal consistency (Cronbach’s alpha). Results evidenced KAP’s domains acceptable internal consistency Cronbach’s alpha of α = 0.75, α = 0.76 and α = 0.76, respectively.

**Reference:**

1. White J & Simon MK. Survey/interview validation rubric for expert panel–VREP; 2014
