## Supplementary material for "Knowledge, attitude and practices on dengue fever among paediatric and adult in-patients in Metro Manila, Philippines": Table S1

**Additional File 2: Profile of pediatric and adult patients with DF and youth and adult controls**

| **Table S1.** Socio-demographic profile, clinical parameters and clinical symptoms among paediatric and adult patients with DF and youth and adult controls | | | | | | | | |
| --- | --- | --- | --- | --- | --- | --- | --- | --- |
|  |  |  | **Patients with DF** | | | **Controls** | | |
|  |  |  | **Total** | **PP** | **AP** | **Total** | **YC** | **AC** |
|  |  | | n = 250 | n = 233 | n = 17 | n = 250 | n = 233 | n = 17 |
| **Socio-Demographic Profile** | Gender | Male | 138 (55.2) | 132 (56.7) | 6 (35.3) | 125 (50.0) | 119 (51.1) | 6 (35.3) |
|  |  | Female | 112 (44.8) | 101 (43.3) | 11 (64.7) | 125 (50.0) | 114 (48.9) | 11 (64.7) |
|  | Age^‡^ | 8-10 | 54 (21.6) | 54 (23.2) |  | 4 (1.60) | 4 (1.70) |  |
|  |  | 11-13 | 53 (21.2) | 53 (22.7) |  | 93 (37.2) | 93 (39.9) |  |
|  |  | 14-16 | 75 (30.0) | 75 (32.2) |  | 111 (44.4) | 111 (47.6) |  |
|  |  | 17-18 | 51 (20.4) | 51 (20.4) |  | 25 (10.0) | 25 (10.7) |  |
|  |  | 19-21 | 5 (2.00) |  | 5 (29.4) | 3 (1.20) |  | 3 (17.6) |
|  |  | 22-24 | 2 (0.80) |  | 2 (11.8) | 4 (1.60) |  | 4 (23.5) |
|  |  | 25-27 | 2 (0.80) |  | 2 (11.8) | 3 (1.20) |  | 3 (17.6) |
|  |  | ≥ 28 | 8 (3.20) |  | 8 (47.1) | 7 (2.80) |  | 7 (41.2) |
|  | Education/ Employment ^‡^ | Grade School | 72 (30.1) | 72 (31.9) |  | 11 (4.40) | 11 (4.70) |  |
|  |  | JHS | 106 (44.4) | 106 (46.9) |  | 204 (81.6) | 204 (87.6) |  |
|  |  | SHS | 39 (16.3) | 36 (15.9) | 3 (23.1) | 12 (4.80) | 12 (5.20) |  |
|  |  | College | 8 (3.30) | 7 (3.10) | 1 (7.70) | 9 (3.60) | 6 (2.60) | 3 (17.6) |
|  |  | Employed | 13 (5.40) | 5 (2.20) | 8 (61.5) | 10 (4.00) |  | 10 (58.8) |
|  |  | Unemployed | 1 (0.40) | 0 (0.00) | 1 (7.70) | 4 (1.60) |  | 4 (23.5) |
|  | Income (₱) | ≤ 10,000 Php | 192 (83.1) | 180 (84.1) | 12 (70.6) | 18 (7.20) | 17 (7.30) | 1 (6.30) |
|  |  | ≥ 10, 000 Php | 39 (16.9) | 34 (15.9) | 5 (29.4) | 231 (92.8) | 216 (92.7) | 15 (93.8) |
|  | Civil Status | Single | 243 (98.4) | 232 (100) | 11 (73.3) | 249 (99.6) | 233 (100) | 16 (94.1) |
|  |  | Married/Live-in | 4 (1.60) | 0 (0.00) | 4 (26.7) | 1 (0.40) | 0 (0.00) | 1 (0.30) |
|  | Household member | ≤ 5 members | 128 (55.4) | 116 (54.2) | 12 (70.6) | 174 (69.6) | 164 (70.4) | 10 (58.8) |
|  |  | ≥ 6 members | 103 (44.6) | 98 (45.8) | 5 (29.4) | 76 (30.4) | 69 (29.6) | 7 (41.2) |
| **Clinical Parameters** | Medical Diagnosis | Dengue w/ ws | 198 (79.2) | 181 (77.7) | 17 (100) |  |  |  |
|  |  | Severe Dengue | 6 (2.40) | 6 (2.57) | 0 (0.00) |  |  |  |
|  |  | Probable | 46 (18.4) | 46 (19.7) | 0 (0.00 |  |  |  |
|  | Days in the Hospital | ≤ 2 days | 170 (76.2) | 159 (77.2) | 11 (64.7) |  |  |  |
|  |  | ≥ 3 days | 53 (23.8) | 47 (22.8) | 6 (35.3) |  |  |  |
|  | DF History | Had DF | 16 (7.30) | 15 (7.3) | 1 (6.70) | 32 (12.8) | 31 (13.3) | 1 (5.90) |
|  |  | First-time | 204 (92.7) | 190 (92.7) | 14 (93.3) | 218 (87.2) | 202 (86.7) | 16 (94.1) |
|  | Family DF History | None | 155 (70.5) | 140 (69.0) | 15 (88.2) | 195 (78.0) | 185 (79.4) | 10 (58.8) |
|  |  | ≥ 1 had DF | 65 (29.5) | 63 (31.0) | 2 (11.8) | 55 (22.0) | 48 (20.6) | 7 (41.2) |
|  | Dengue Phase | Acute | 200 (80.0) | 188 (80.7) | 12 (70.6) |  |  |  |
|  |  | Recovery | 50 (20.0) | 45 (19.3) | 5 (29.4) |  |  |  |
|  | Dengue Tests | (-) NS1Ag | 52 (44.1) | 42 (38.9) | 10 (100) |  |  |  |
|  |  | (+) NS1Ag | 66 (55.9) | 66 (61.1) | 0 (0.00) |  |  |  |
|  |  | (-) IgG | 46 (68.7) | 41 (67.2) | 5 (83.3) |  |  |  |
|  |  | (+) IgG | 21 (31.3) | 20 (32.8) | 1 (16.7) |  |  |  |
|  |  | (-) IgM | 14 (20.9) | 13 (21.3) | 1 (16.7) |  |  |  |
|  |  | (+) IgM | 53 (79.1) | 48 (78.7) | 5 (83.3) |  |  |  |
| **Clinical Symptoms** | Headache | Asymptomatic | 206 (82.4) | 193 (82.8) | 13 (76.5) |  |  |  |
|  |  | Symptomatic | 44 (17.6) | 40 (17.3) | 4 (23.5) |  |  |  |
|  | Fever | Asymptomatic | 210 (84.0) | 194 (83.3) | 16 (94.1) |  |  |  |
|  |  | Symptomatic | 40 (16.0) | 39 (16.9) | 1 (5.9) |  |  |  |
|  | Nausea & Vomiting | Asymptomatic | 188 (75.2) | 177 (76.0) | 11 (64.7) |  |  |  |
|  |  | Symptomatic | 62 (24.8) | 56 (24.3) | 6 (35.3) |  |  |  |
|  | Myalgias & Arthralgias | Asymptomatic | 193 (77.2) | 180 (77.3) | 13 (76.5) |  |  |  |
|  |  | Symptomatic | 57 (22.8) | 53 (22.9) | 4 (23.5) |  |  |  |
|  | Petechiae (Rash) | Asymptomatic | 145 (58.0) | 134 (57.5) | 11 (64.7) |  |  |  |
|  |  | Symptomatic | 105 (42.5) | 99 (43.0) | 6 (35.3) |  |  |  |
|  | Retro-/ Peri-Orbital Pain | Asymptomatic | 223 (89.2) | 208 (89.3) | 15 (88.2) |  |  |  |
|  |  | Symptomatic | 27 (10.8) | 25 (7.86) | 2 (11.8) |  |  |  |
|  | Abdominal Pain | Asymptomatic | 157 (62.8) | 145 (62.2) | 12 (70.6) |  |  |  |
|  |  | Symptomatic | 93 (37.2) | 88 (37.9) | 5 (29.4) |  |  |  |
|  | Thrombocytopenia | ≤ 9900/mm3 | 72 (28.8) | 70 (22.01) | 2 (11.8) |  |  |  |
|  |  | ≥ 10000/mm3 | 165 (66.0) | 150 (68.2) | 15 (88.2) |  |  |  |
| PP = Paediatric Patients; AP = Adult Patients; YC = Youth Controls; AC = Adult Controls; ₱ = Philippine Peso (52.16 USD = 1 ₱); Acute = febrile to critical phase; WS = warning signs; DF = Dengue Fever; (+) = positive; (-) = negative; NS1Ag = Non-Structural protein 1 antigen; IgG = Immunoglobin G antibody; IgM = Immunoglobin M antibody | | | | | | | | |
