## Supplementary material for "Knowledge, attitude and practices on dengue fever among paediatric and adult in-patients in Metro Manila, Philippines": Table S2

**Additional file 3 – Detailed information on the sources of information on dengue fever of pediatric and adult patients with DF and controls**

| **Table S2.** Sources of information on dengue fever of paediatric and adult patients with DF and youth and adult controls | | | | | | | | | | | | | | | | | | | | | | | | | | | | | | | | | | | |
| --- | --- | --- | --- | --- | --- | --- | --- | --- | --- | --- | --- | --- | --- | --- | --- | --- | --- | --- | --- | --- | --- | --- | --- | --- | --- | --- | --- | --- | --- | --- | --- | --- | --- | --- | --- |
|  |  | **Patients with DF** | | |  | **Controls** | | |  | **PP vs YC** | **AP vs AC** | **Knowledge** | | | | | | | | **Attitude** | | | | | | | | **Practice** | | | | | | | |
|  |  | **Total** | **Paediatric** | **Adults** |  | **Total** | **Youth** | **Adults** |  |  |  | **PP** | | **AP** | | **YC** |  | **AC** |  | **PP** | | **AP** | | **YC** |  | **AC** |  | **PP** | | **AP** | | **YC** |  | **AC** |  |
|  | | n = 250 | n = 233 | n = 17 |  | n = 250 | n = 233 | n = 17 |  |  |  | **Mean** | | **Mean** | | **Mean** | | **Mean** | | **Mean** | | **Mean** | | **Mean** | | **Mean** | | **Mean** | | **Mean** | | **Mean** | | **Mean** | |
| Social Media | | | | | |  |  |  |  | 0.01 | 0.73 |  |  |  |  |  |  |  |  |  |  |  |  |  |  |  |  |  |  |  |  |  |  |  |  |
|  | No | 129 (51.6) | 122 (52.4) | 7 (41.2) |  | 100 (40.0) | 92 (39.5) | 8 (47.1) |  |  |  | 19.3 |  | 15.7 |  | 13.5 | * | 23.2 |  | 3.32 |  | 3.28 |  | 2.42 |  | 3.75 |  | 9.07 |  | 6.86 |  | 9.76 |  | 10.7 |  |
|  | Yes | 121 (48.4) | 111 (47.6) | 10 (58.8) |  | 150 (60.0) | 141 (60.5) | 9 (52.9) |  |  |  | 20.0 |  | 19.1 |  | **15.1** |  | 21.8 |  | 3.67 |  | 4.30 |  | 2.49 |  | 4.44 |  | 9.52 |  | 8.70 |  | 9.88 |  | 11.3 |  |
| TV | | | | | |  |  |  |  | 0.32 | 0.22^a^ |  |  |  |  |  |  |  |  |  |  |  |  |  |  |  |  |  |  |  |  |  |  |  |  |
|  | No | 62 (24.8) | 60 (25.7) | 2 (11.8) | ^a^ | 70 (28.0) | 64 (27.5) | 6 (35.3) | ^a^ |  |  | 19.3 |  | 18.0 |  | 14.0 |  | 22.7 |  | 3.21 |  | 6.00 |  | 2.62 |  | 3.83 |  | 9.30 |  | 7.00 |  | 9.56 |  | 11.2 |  |
|  | Yes | 188 (75.2) | 173 (74.2) | 15 (88.2) |  | 180 (72.0) | 169 (72.5) | 11 (64.7) |  |  |  | 19.8 |  | 17.7 |  | 14.6 |  | 22.4 |  | 3.58 |  | 3.60 |  | 2.40 |  | 4.27 |  | 9.31 |  | 8.07 |  | 9.93 |  | 11.0 |  |
| Radio | | | | | |  |  |  |  | 0.001 | 0.16 |  |  |  |  |  |  |  |  |  |  |  |  |  |  |  |  |  |  |  |  |  |  |  |  |
|  | No | 131 (52.4) | 123 (52.8) | 8 (47.1) |  | 169 (67.6) | 157 (67.4) | 12 (70.6) |  |  |  | 19.4 |  | 17.0 |  | 14.3 |  | 23.1 |  | 3.34 |  | 3.12 |  | 2.48 |  | 3.75 |  | 9.09 |  | 8.00 |  | 9.64 |  | 10.7 |  |
|  | Yes | 119 (47.6) | 110 (47.2) | 9 (52.9) |  | 81 (32.4) | 76 (32.6) | 5 (29.4) |  |  |  | 19.9 |  | 18.3 |  | 14.8 |  | 21.0 |  | 3.65 |  | 4.55 |  | 2.43 |  | 5.00 |  | 9.50 |  | 7.89 |  | 10.2 |  | 11.8 |  |
| Newspapers | | | | | |  |  |  |  | < 0.001 | 0.33^a^ |  |  |  |  |  |  |  |  |  |  |  |  |  |  |  |  |  |  |  |  |  |  |  |  |
|  | No | 165 (66.0) | 149 (63.9) | 16 (94.1) | * | 203 (81.2) | 190 (81.5) | 13 (76.5) | ^a^ |  |  | 19.1 | ** | 17.43 |  | 13.7 | *** | 23.1 |  | 3.26 | * | 3.75 |  | 2.33 | * | 3.92 |  | 8.93 | ** | 8.00 |  | 9.61 | ** | 10.6 |  |
|  | Yes | 85 (34.0) | 84 (36.0) | 1 (5.9) |  | 47 (18.8) | 43 (18.5) | 4 (23.5) |  |  |  | **20.5** |  | 22.0 |  | **17.7** |  | 20.5 |  | **3.89** |  | 6.00 |  | **3.07** |  | 4.75 |  | **9.90** |  | 7.00 |  | **10.8** |  | 12.5 |  |
| Health brochures | | | | | |  |  |  |  | < 0.001 | 0.49 |  |  |  |  |  |  |  |  |  |  |  |  |  |  |  |  |  |  |  |  |  |  |  |  |
|  | No | 162 (64.8) | 153 (65.7) | 9 (52.9) |  | 93 (37.2) | 194 (83.3) | 11 (64.7) | ^a^ |  |  | 19.4 |  | 17.3 |  | 13.9 | ** | 22.7 |  | 3.22 | ** | 3.33 |  | 2.35 |  | 4.00 |  | 9.25 |  | 8.11 |  | 9.69 | * | 11.3 |  |
|  | Yes | 88 (35.2) | 80 (34.3) | 8 (47.1) |  | 45 (18.0) | 39 (16.7) | 6 (35.3) |  |  |  | 20.1 |  | 18.1 |  | **17.2** |  | 22.0 |  | **3.99** |  | 4.50 |  | 3.02 |  | 4.33 |  | 9.35 |  | 7.75 |  | **10.5** |  | 10.7 |  |
| Family | | | | | |  |  |  |  | < 0.001 | 0.30 |  |  |  |  |  |  |  |  |  |  |  |  |  |  |  |  |  |  |  |  |  |  |  |  |
|  | No | 145 (58.0) | 135 (57.9) | 10 (58.8) |  | 93 (37.2) | 86 (36.9) | 7 (41.2) |  |  |  | 19.4 |  | 17.3 |  | 13.0 | * | 23.7 |  | 3.48 |  | 3.10 |  | 2.17 |  | 4.00 |  | 9.10 |  | 7.50 |  | 9.12 | ** | 11.1 |  |
|  | Yes | 105 (42.0) | 98 (42.1) | 7 (41.2) |  | 157 (62.8) | 147 (63.1) | 10 (58.8) |  |  |  | 20.0 | .0 | 18.3 |  | **15.3** |  | 21.6 |  | 3.50 |  | 5.00 |  | 2.62 |  | 4.20 |  | 9.54 |  | 8.57 |  | **10.2** |  | 11.0 |  |
| Neighbours | | | | | |  |  |  |  | < 0.001 | 0.45 |  |  |  |  |  |  |  |  |  |  |  |  |  |  |  |  |  |  |  |  |  |  |  |  |
|  | No | 163 (65.2) | 152 (65.2) | 11 (64.7) |  | 219 (87.6) | 206 (88.4) | 13 (76.5) | ^a^ |  |  | 19.8 |  | 18.1 |  | 14.4 |  | 22.6 |  | 3.44 |  | 3.64 |  | 2.38 |  | 3.61 | ** | 9.31 |  | 8.18 |  | 9.75 |  | 11.0 |  |
|  | Yes | 87 (34.8) | 81 (34.8) | 6 (35.3) |  | 31 (12.4) | 27 (11.6) | 4 (23.5) |  |  |  | 19.4 |  | 17.0 |  | 14.6 |  | 22.0 |  | 3.59 |  | 4.33 |  | 3.07 |  | **5.75** |  | 9.25 |  | 7.50 |  | 10.4 |  | 11.2 |  |
| School | | | | | |  |  |  |  | 0.04 | 1.00 |  |  |  |  |  |  |  |  |  |  |  |  |  |  |  |  |  |  |  |  |  |  |  |  |
|  | No | 136 (54.4) | 128 (54.9) | 8 (47.1) |  | 111 (44.4) | 103 (44.2) | 8 (47.1) |  |  |  | 19.4 |  | 17.0 |  | 12.1 | *** | 22.6 |  | 3.28 |  | 2.75 |  | 1.89 | *** | 3.62 |  | 9.27 |  | 7.75 |  | 9.22 | ** | 10.6 |  |
|  | Yes | 114 (45.6) | 105 (45.1) | 9 (52.9) |  | 139 (55.6) | 130 (55.8) | 9 (52.9) |  |  |  | 19.9 |  | 18.3 |  | **16.4** |  | 22.3 |  | 3.74 |  | 4.89 |  | **2.91** |  | 4.55 |  | 9.28 |  | 8.11 |  | **10.3** |  | 11.4 |  |
| Hospital, Doctors, Nurses | | | | | |  |  |  |  | < 0.001 | 0.006 |  |  |  |  |  |  |  |  |  |  |  |  |  |  |  |  |  |  |  |  |  |  |  |  |
|  | No | 78 (31.2) | 73 (31.3) | 5 (29.4) |  | 130 (52.0) | 117 (50.2) | 13 (76.5) |  |  |  | 19.4 |  | 17.0 |  | 12.8 | *** | 23.1 |  | 2.90 | ** | 2.80 |  | 2.16 | * | 3.92 |  | 9.50 |  | 7.80 |  | 9.00 | *** | 10.7 |  |
|  | Yes | 172 (68.8) | 160 (70.2) | 12 (70.6) |  | 120 (48.0) | 116 (49.8) | 4 (23.5) |  |  |  | 19.7 |  | 18.0 |  | **16.1** |  | 20.5 |  | **3.74** |  | 4.33 |  | **2.76** |  | 4.75 |  | 9.20 |  | 8.00 |  | **10.7** |  | 12.2 |  |
| Barangay and community | | | | | |  |  |  |  | < 0.001 | 0.13 |  |  |  |  |  |  |  |  |  |  |  |  |  |  |  |  |  |  |  |  |  |  |  |  |
|  | No | 147 (58.8) | 137 (58.8) | 10 (58.8) |  | 215 (86.0) | 201 (86.3) | 8 (47.1) | ^a^ |  |  | 19.6 |  | 17.7 |  | 14.3 |  | 22.5 |  | 3.23 | * | 3.30 |  | 2.39 |  | 4.00 |  | 9.15 |  | 8.00 |  | 9.66 | * | 10.6 | * |
|  | Yes | 103 (41.2) | 96 (41.2) | 7 (41.2) |  | 35 (14.0) | 32 (13.7) | 3 (17.6) |  |  |  | 19.7 |  | 17.7 |  | 15.7 |  | 22.3 |  | **3.85** |  | 4.71 |  | 2.90 |  | 6.67 |  | 9.48 |  | 7.86 |  | **10.9** |  | **13.0** |  |
| Workplace | | | | | |  |  |  |  | < 0.001 | 0.17^a^ |  |  |  |  |  |  |  |  |  |  |  |  |  |  |  |  |  |  |  |  |  |  |  |  |
|  | No | 206 (82.4) | 194 (83.3) | 12 (70.6) | ^a^ | 239 (95.6) | 223 (95.7) | 16 (94.1) | ^a^ |  |  | 19.6 |  | 17.4 |  | 14.4 |  | 22.5 |  | 3.42 |  | 3.50 |  | 2.47 |  | 4.00 |  | 9.26 |  | 7.58 |  | 9.77 |  | 10.8 | * |
|  | Yes | 44 (17.6) | 39 (16.7) | 5 (29.4) |  | 11 (4.4) | 10 (4.3) | 1 (5.9) |  |  |  | 20.0 |  | 18.4 |  | 15.7 |  | 22.0 |  | 3.90 |  | 4.80 |  | 2.30 |  | 6.00 |  | 9.36 |  | 8.80 |  | 11.2 |  | **14.0** |  |
| Health centre | | | | |  |  |  |  |  | < 0.001 | 0.002 |  |  |  |  |  |  |  |  |  |  |  |  |  |  |  |  |  |  |  |  |  |  |  |  |
|  | No | 90 (36.0) | 194 (36.5) | 5 (29.4) |  | 191 (76.4) | 177 (76.0) | 14 (82.4) | ^a^ |  |  | 19.9 |  | 14.2 | ** | 14.0 | * | 22.9 |  | 3.13 | * | 2.80 |  | 2.40 |  | 3.86 |  | 9.09 |  | 8.20 |  | 9.47 | ** | 10.6 |  |
|  | Yes | 160 (64.0) | 148 (63.5) | 12 (70.6) |  | 59 (23.6) | 56 (24.0) | 3 (17.6) |  |  |  | 19.5 |  | **19.2** |  | **16.1** |  | 20.3 |  | **3.68** |  | 4.33 |  | 2.66 |  | 5.33 |  | 9.40 |  | 7.83 |  | 11.0 |  | 13.3 |  |
| PP = Paediatric Patients; AP = Adult Patients; YC = Youth Controls; AC = Adult Controls a = p-values based on Fisher’s exact test; * P < 0.05, ** P < 0.01, ***P < 0.001 | | | | | | | | | | | | | | | | | | | | | | | | | | | | | | | | | | | |
